# A network based study of the dynamics of *Aβ* and τ proteins in Alzheimer’s disease

**DOI:** 10.1101/2024.10.17.618808

**Authors:** Stefano Bianchi, Germana Landi, Camilla Marella, Maria Carla Tesi, Claudia Testa, Alzheimers Disease Neuroimaging Initiative

**Author notes:** Data used in preparation of this article were obtained from the Alzheimers Disease Neuroimaging Initiative (ADNI) database (adni.loni.usc.edu). As such, the investigators within the ADNI contributed to the design and implementation of ADNI and/or provided data but did not participate in analysis or writing of this report. A complete listing of ADNI investigators can be found at: http://adni.loni.usc.edu/wp-content/uploads/how_to_apply/ADNI_Acknowledgement_List.pdf. These authors contributed equally to this work.

## Abstract

Due to the extreme complexity of Alzheimer’s disease (AD), the aetiology of which is not yet known, nor are there any known effective treatments, mathematical modelling can be very useful. Indeed, mathematical models, if deemed reliable, can be used to test medical hypotheses that could be difficult to verify directly. In this context, it is important to understand how *Aβ* and *τ* proteins, which in abnormal aggregate conformations are hallmarks of the disease, interact and spread. We are particularly interested in this paper in studying the spreading of misfolded *τ* . To this end, we present four different mathematical models, all on networks on which the protein evolves. The models differ in both the choice of network and diffusion operator. Through comparison with clinical data on *τ* concentration, that we carefully obtained with multimodal analysis techniques, we show that some models are more adequate than others to simulate the dynamics of the protein. This type of study may suggest that, when it comes to modelling certain pathologies, the choice of the mathematical setting must be made with great care if comparison with clinical data is considered decisive.

## 1 Introduction

Alzheimers disease (AD) is a neurodegenerative disorder characterized by a progressive decline in memory and other cognitive functions, leading inevitably to death. It is an incurable disease affecting more than 50 million people, with figures set to increase significantly in the coming years (World Alzheimer Report 2023). The aetiology of the disease is unclear to date, but two proteins are universally recognised to play a crucial role in the development of the disease: amyloid-beta (*Aβ*) and tau (*τ*). Both proteins are physiologically present in the brain, but in the presence of the disease they form abnormal aggregates, in a progressive and irreversible way. Indeed *Aβ* plaques and neurofibrillary tangles (NFT) of *τ* are pathological hallmarks of AD. Experimental studies suggest non-uniform distributions of pathological proteins in the brain [32]. Both *Aβ* and *τ* exhibit characteristic spatiotemporal deposition patterns. NFT tangles appear first in the entorhinal cortex and then spread to the amygdala, temporal areas and lately throughout the cortex [7]. On the other hand, *Aβ* plaques form first in the temporal and frontal areas and then spread to other areas of the brain [7, 20]. Recent literature suggests that the interplay between the two proteins should be crucial in the development of the disease and must be taken into account for the development of new therapies [9, 45, 4]. Clearly when one is interested in modelling complex pathologies such as AD, comparison with clinical and experimental data is crucial to the reliability of the study. Clinical multimodal neuroimages, if properly processed in terms of harmonization, combination and quantitative approach can give information of local character, providing for instance the concentration of proteins in different brain regions [25].

This work has two main objectives. One is to capture such types of non-uniform distributions through appropriate mathematical models. The other purpose is to compare the results from the mathematical models with clinical data in the context of Alzheimer’s disease. Comparison with clinical data is a fundamental step in the development of a trustable mathematical model but it is rarely addressed in the mathematical modelling literature. To this aim, we have processed a considerable amount of clinical neuroimages to extract quantitative features which can be compared with the *τ* concentration values provided by the mathematical models.

Networks are mathematical tools which provide an ideal setting for comparisons between clinical data and numerical results from models. Complex network properties have been identified with some consistency in all modalities of neuroimaging data and over a range of spatial and time scales. Thus, brain networks allow to describe the information process in neurons and its characteristic of being both locally specialized or segregated and globally distributed or integrated [2]. In particular, small-worldness characterizes an intermediate regime of networks between the two extreme regimes of a regular lattice and a random network, efficiently representing the brain structure [41]. Mathematical modeling of AD can be very useful for integrating clinical and experimental data in a theoretical setting which could allow, once reliability has been verified, to test various hypotheses otherwise difficult to verify. For an exhaustive overview of existing mathematical models of proteins spreading on networks and of various related challenging questions we refer to [46]. Here we wish to emphasise that the dynamics of the two proteins we are interested in differ in several respects, both spatial (*Aβ* spreads over small distances, *τ* over large distances) and temporal (the dynamics of *Aβ* is much faster than that of *τ*). Differences in spatial dynamics should in some way be reflected by the choice of network used for the evolution of the dynamics. This paper is inspired by the paper [5] as far as the idea of using distinct networks for different protein dynamics is concerned; however its purpose is quite different. Indeed, in [5] the authors were mainly interested in investigating the synergisitic interactions of the two proteins *Aβ* and *τ*, whose relevance was confirmed there by testing various modelling hypotheses. However the authors were not interested to any comparison with medical data. Here we strongly rely on the comparison with medical data to test different modelling hypothesis concerning diffusion of *τ* on properly chosen networks, to see if there is an optimal way to deal with the spreading of proteins when it comes to modelling AD. Therefore the main target of this paper is twofold: 1. to consider different models for the spreading of *Aβ* and especially of *τ*, developed on networks with a precise biomedical meaning; 2. to have medical data for a comparison as meaningful as possible with the models, and use that comparison to test the trustability of the model.

The plan of the work is the following: in Section 2 we describe the networks we use for our models. We start from data publicly available, and from that we construct several different weighted graphs on which we evolve proteins. In Section 3 we describe the models we study: in particular, we present four different models for various possible ways of the spreading of *τ* . Section 4 is concerned with the procedures we have adopted to obtain reliable medical data. We emphasize that we used a multimodal approach, based on MRI and PET, to determine the concentrations of *τ* protein in the brain. Finally in Section 5 we present the results obtained by numerical simulations and we make comparisons with medical data.

## 2 Brain networks via weigheted graphs

The central concept of a brain network constructed to segregate and integrate information processing comes from the advent of ”disconnection syndromes” hypothesis [10] mainly on the basis of clinico-pathological correlations. The conceptualization of the relationship between structure and function of the brain has lead to understand that brain regions participate to many functions introducing the structure-function framework described by a network of brain regions. From Mesulam and over [27], it has been introduced the network approach to understand the localization of complex functions. The connection matrix of the human brain, known as the human ”connectome”, today represents an indispensable tool for mapping brain structure to functional processes and has a valuable impact on understanding brain diseases.

To model the spreading of different proteins in the brain, we consider several connectomes, each corresponding to a different undirected weighted graph. Data used for the construction of these connectomes have been downloaded at the website https://braingraph.org and consist of an *averaged* graph of 477 healthy subjects each with 1015 nodes (the weight calculation mode set as median and the number of fibers launched set to 20) [43, 42]. We stress that the vertices of such a graph do not have associated coordinates that determine their position in space, since it is precisely an averaged graph. This will have consequences when we talk about “distances”, which must be understood in an intrinsic sense and not in the classical euclidean metric sense.

In the following, we first recall the definition of weighted graph and how to construct an associated Laplacian, mimicking diffusion on it. Then, we describe the weighted graphs we will use to model the spreading of *Aβ* and *τ* proteins. A *graph* is a pair 𝒢 = (*V, E*), where *V* is a set of vertices and *E* ⊂ *V* × *V* is a set of the edges; 𝒢 is said to be undirected if (*i, j*) ∈ *E* implies that (*j, i*) ∈ *E*. The graph G is called weighted if a measure *ω* : *E* → ℝ^+^ exists, assigning a unique positive number to each edge; the value *ω*(*i, j*) is referred to as weight of the edge (*i, j*). A weighted graph can be represented through the adjacency matrix *A* whose entries *A*_*i*,*j*_ represent the weights of the edge (*i, j*). Let *N* = |*V* | be the number of vertices in 𝒢, then *A* ∈ R^*N×N*^ is defined as

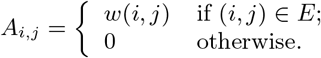

There are several possible definitions for the Laplacian associated to a given graph. Given the adjacency matrix *A* ∈ ℝ^*N×N*^ of 𝒢, following [19] we define the graph Laplacian *L* as *L* = *D* − *A* where *D* is the weighted degree matrix whose *j*th diagonal element *D*_*j*,*j*_ is given by

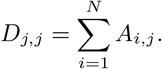

### 2.1 Structural Connectome

We call *structural connectome* a weighted graph 𝒢 = (*V, E*) extracted from tractography of diffusion tensor images of 477 healthy subjects of the Human Connectome Project [26] using the Budapest Reference Connectome v3.0 [42]. In this graph, vertices correspond to parcellated regions of grey matter formed by neurons which share similarity in cytoarchitecture, functional activity, and structural connections to other regions, and edges represent the connectivity between the regions. We use a high-resolution connectome with *N* = 1015, which can be downloaded at the website https://pitgroup.org/connectome/. It is possible to choose for the edges of this graph different weights, based on the mean number of fibers connecting two regions, and on their mean lenght. Accordingly, we will denote by 𝒢_NL_ the graph with weights given by

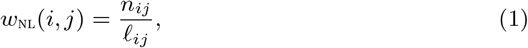

where *n*_*ij*_ is the mean number of fibers connecting verices *i* and *j*, and *R*_*ij*_ is the mean lenght of such fibers. Analogously, we will denote by 𝒢_l_ the graph with weights given by

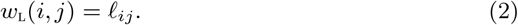

Therefore, the two weighetd graphs 𝒢_NL_ and 𝒢_L_ have the same vertices and edges, but different weights.

### 2.2 Intrinsic Proximity Connectome

Let *R*_P_ ∈ ℝ^+^ be a given positive value, the *intrinsic proximity connectome* is the weighted graph 𝒢_P_ = (*V, E*) whose set of vertices *V* is the same as 𝒢_L_’s and whose set of edges *E*_P_ ⊂ *E*_L_ is the subset of edges of 𝒢_L_ connected by a fiber with length *𝓁* less than *R*_P_. The weights are given by

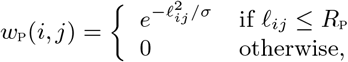

where *σ* ∈ ℝ^+^ is a fixed parameter. In this way, we connect two vertices only if they are close in an intrinsic sense, as opposed to a “geometric” vicinity measured with some kind of euclidean distance. Indeed, as already mentioned the vertices of the graph do not have associated spatial coordinates. The weights we assign are stronger for intrinsically close vertices. For this reason, 𝒢_p_ is referred to as *intrinsic proximity connectome*. It is appropriate to emphasize here that this proximity connectome is quite different from the one used in [5]. Indeed here we use only *intrisic* connections, without adding any new connection, and we give them a weight proportional to the intrinsic distance.

### 2.3 Cumulative Connectome

The *cumulative connectome* is a weighted graph 𝒢_C_ = (*V, E*) with the same *V* and *E* as 𝒢_NL_, but with weights defined as follows. Let 𝒮_*ij*_ be the set of all paths in the graph 𝒢_NL_ starting at vertex *i* and ending at vertex *j*, i.e:

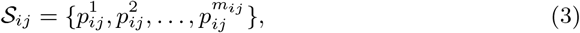

where *m*_*ij*_ = |𝒮 _*ij*_ | and each path 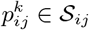 consists of a sequence of edges:

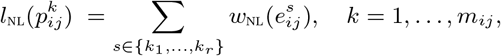

Recall that the graphs 𝒢_NL_ and 𝒢 _L_ differ only in the weights assigned to the edges, thus we can define the length of the path 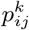 considering it living in the graph 𝒢_l_ or in the graph 𝒢_NL_. Since the length of a path is usually defined as the sum of the weights of the edges forming the path, in 𝒢_NL_ we obtain the length 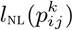 defined as

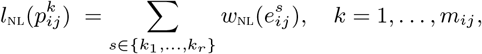

and, in 𝒢_L_we obtain the length 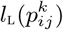 given by

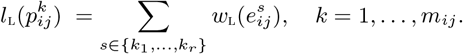

Finally let 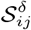 be the subset of 𝒮 such that

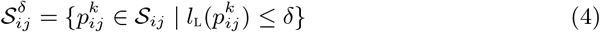

where *δ* ∈ ℝ^+^ is a fixed parameter. Two vertices *i* and *j* are connected in the cumulative connectome 𝒢_C_ if there is at least one path of length in 𝒢_L_ less or equal *δ* joining them (we call such a path an admissible path.). In the affirmative case, the weight of the edge connecting vertices *i* and *j* is given by the sum of the lengths in 𝒢_NL_ of all admissible paths (hence the name cumulative connectome):

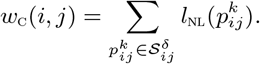

Summarising, in the cumulative connectome, two vertices *i* and *j* are connected if there is at least one path starting at *i* and ending at *j* whose length is smaller than a fixed value *δ*. Vertices that are not connected in 𝒢_NL_ will be connected in 𝒢_C_ if there is an admissible path in 𝒢_NL_ joining them. In this way, we encode in 𝒢 _c_ connections between brain regions over long distances. In other words, we connect two vertices if the corresponding brain regions are joined by axonal paths with length determined by a *δ* parameter that we choose. The weights of the connections depend on how many and how long the axonal fibers in the connections are.

## 3 The models

By choosing appropriate graphs among the ones introduced in the previous section, we set up different models to describe the dynamics of *Aβ* and *τ* proteins. The choice of the graphs is guided by the main biological features characterizing the two proteins and their dynamics.

Concerning *Aβ*, it is well known that monomeric *Aβ* peptides originate physiologically from the cleavage of the transmembrane protein APP (amyloid precursor protein), and are considered not toxic [16]. Then, for reason not yet clarified, an imbalance between production and clearence of the protein can occur, giving rise to a process of agglomerations. This leads to the formation of toxic amyloid fibrils [22, 31], often referred to as olygomers. These olygomers eventually accumulate in insoluble agglomerates known as senile plaques, nowadays considered not toxic [22]. To describe this process we adopt a so-called compartmental model, that is we consider *Aβ* proteins assuming only three types of conformations: monomers, olygomers (which include all soluble conformations that are not monomers) and plaques (insoluble conformations).

Moreover, we want to take into account that *Aβ* protein diffuses on short distances and its dynamics (production, aggregation, clearence) is fast [29]. The molar concentration of a protein on the vertices *V* of a graph will be denoted by an *N* -dimensional vector, where *N* = |*V* | is the number of vertices of the graph. Accordingly, the vector valued functions **u**_*j*_ : ℝ → ℝ^*N*^ describe the molar concentration of *Aβ* monomers (*j* = 1), olygomers (*j* = 2) and plaques (*j* = 3).

We model *Aβ* dynamics with the following equations [5]:

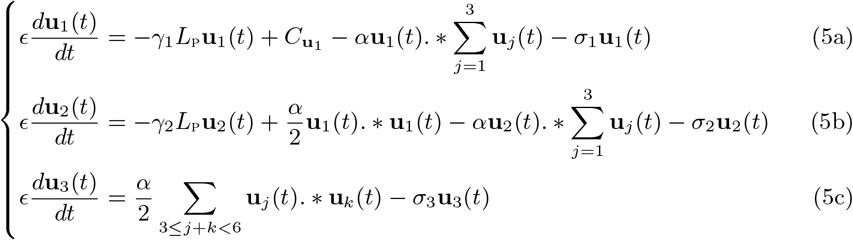

where .* denotes the element-wise product between vectors. System (5) is endowed with initial conditions at *t* = 0 as follows (at the initial time the brain is healthy, therefore only *Aβ* monomers are present):

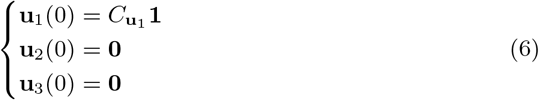

where **0** and **1** denote the *N* -dimensional vectors whose components are all equal to zero and one, respectively. Since *t* denotes a slow time variable, the *E* in front of the equations takes into account the fact that the processes described are fast [29]. The first term in equations (5a) and (5b) models diffusion of *Aβ* monomers and olygomers along the network. Since this protein spreads along short distances, we choose the intrinsic proximity connectome to model the network. Indeed on the intrinsic proximity connectome 𝒢 _p_ two vertices *i* and *j* are connected only if the mean length of the axonal fibers connecting the corresponding regions of the brain is sufficiently short; in addition the shorter the fibre length, the greater the weight of the corresponding edge. Accordingly, here diffusion is driven by the graph Laplacian *L*_p_. In the first equation, the term 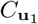 represents a source term (gain) due to the physiological production of *Aβ* monomers, the term - *α***u**_1_(*t*).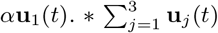 a loss due to the aggregtion of monomers with other *Aβ* proteins and the term −*σ*_1_**u**_1_(*t*) a loss due to clearence phenomena. In the second equation the term 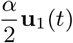. * **u** (*t*) is a gain in olygomers due to the aggregation of two monomers, while the next two term are losses analogous to those in the first equation. Finally, in the third equation the diffusion term is absent since plaques are insoluble (they are too heavy to diffuse), there is a gain term due to aggregation (for a detailed explanation of this term and in general on the use of Smoluchowki’s equations in this context see [4]) and a loss term due to clearence. We stress that in the equations governing the dynamics of *Aβ* proteins there is no coupling with the *τ* protein. Indeed, such a coupling will be present in the equation governing the dynamics of *τ* .

Protein *τ* is a physiological microtubule-associated protein: its main function is to stabilise microtubules and to regulate axonal transport. In brains affected by AD it has been found misfolded *τ* in form of aggregates called neurofribrillary tangles, which together with *Aβ* plaques are an hallmark of AD [21]. Misfolded *τ* is toxic for the neuron: indeed it compromises stabilization, transport and in general interferes with neuronal functions. As already mentioned in the Introduction, in recent years there has been a consensus in the scientific community to consider a synergistic effect of *Aβ* and *τ* when it comes to AD [24, 38, 9]. More precisely, the “trigger and bullet” hypothesis identifies in toxic *Aβ* peptides the trigger for *τ* misfolding: the bullet is the misfolded *τ* which in turn causes neural damage until eventually neuron’s death [6, 3, 40]. Concerning the spreading mechanism of misfolded *τ*, it has been proposed that neuronal damage spreads in the neuronal net through a neuron-to-neuron prion-like propagation mechanism [8, 17, 44, 12]. Based on the considerations made so far, the dynamics of misfolded *τ*, whose concentration is given by the vector valued function **w** : ℝ → ℝ^*N*^, is governed by the following equation [5]:

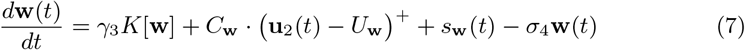

endowed with the initial condition (at the initial time the brain is healthy, therefore there is no misfolded *τ*):

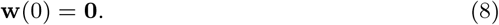

Here we see the coupling between *Aβ* and *τ* . Indeed the second term *C*_**w**_ ·(**u**_2_(*t*)−*U*_**w**_) in the equation rules the interaction of the two proteins: if **u**_2_, the toxic form of *Aβ*, is above a given threshold, then it triggers the misfolding of *τ* . The term *s*_**w**_ (*t*) represents a source of misfolded *τ*, tipically located in the enthorinal region of the brain [7], while the last term is the loss due to clearence.

The main purpose of this work is to investigate different possible forms for the operator *K*, appearing as first term in (7), modelling the spreding of *τ* . As already mentioned, the spreading of such protein can take place over long distances [1], possibly following a prion-like type of process [44], on a slow time scale. Several models have appeared in literature concerning various possible mechanisms of *τ* spreading on networks [36, 37, 35], [33, 34],[14, 15, 18], [5]. Here we consider four different possibilities (corresponding to four different mathematical models) for the operator *K*, and we make a comparison between the results obtained for each model and clinical data. In this way we try to determine if there is one form for *K* better describing the dynamics we are interested in reproducing. The full system of equations we are going to study is therefore given by:

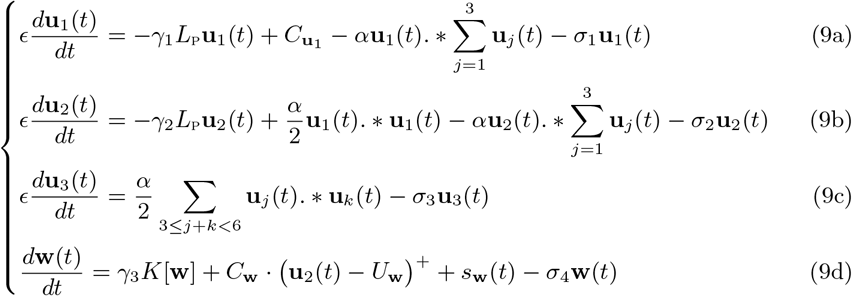

### 3.1 Model 1: diffusion of *τ* along the structural connectome *𝒢*_nl_

Following previous works in literature [36, 14][5, 4], a quite standard choice to model the spreading of *τ* protein along edges of a connectome consists in choosing the Laplacian *L*_NL_ associated to the structural connectome 𝒢_NL_. In this way spreading corresponds to diffusion and we have

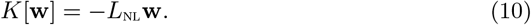

### 3.2 Model 2: diffusion of *τ* along the cumulative connectome 𝒢_c_

In the cumulative connectome, two vertices are connected if there exists at least one admissible path between them. Such a path can be formed by several consecutive edges, each with its own weight contributing to the final weight. The corresponding Laplacian *L*_C_ therefore encodes the structure of the brain with connections spanning long distances. For this reason, we choose

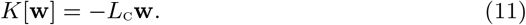

We stress again that all distances mentioned in the paper are to be understood not in a geometric but in an intrinsic sense, i.e. referring to fibre lengths, that is edges, in the graph.

### 3.3 Model 3: spreading of *τ* via convolution on 𝒢_l_

A common way of dealing with dynamics occurring over long distances is to make use of integral operators, tipically convolutions with an appropriate kernel taking into account distances. With notation similar to (4), let 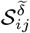 be the subset of 𝒮_*ij*_ containing all the paths in **𝒢**_L_ from node *i* to node *j* with length less than 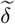:

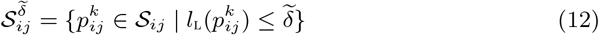

where 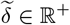 is a fixed parameter. We define the signal *k*_*i*_ on each node *i* of the graph 𝒢_L_ as follows:

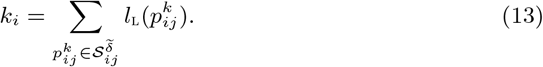

Intuitively, *k*_*i*_ accounts for all the paths between *i* and each vertex *j* in 𝒢_L_ that is not too far from *i* and plays the role of the convolution kernel. The convolution product on 𝒢_L_ between **k** and **w** is defined by using the graph Laplacian eigenvectors [39]. Let {**v**_.*ℓ*_}_*ℓ*=1,…,*N*_ be a complete set of orthonormal eigenvectors of the Laplacian *L*_L_ of 𝒢_L_ with eigenvalues {*λ*_*ℓ*_}_. *ℓ*=1,…,*N*_ ordered in nondecreasing order and let *U* ∈ ℝ^*N×N*^ be the matrix whose columns correspond to the eigenvectors **v**_*ℓ*_, *ℓ* = 1, …, *N* . The graph Fourier transform 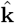 of the signal **k** on 𝒢 _L_ is defined as the expansion of **k** in terms of the eigenvectors of *L*_L_, i.e.:

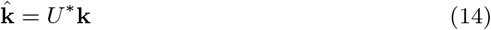

where ^*\**^ denotes the conjugate transpose of a matrix. The inverse graph Fourier transform is given by

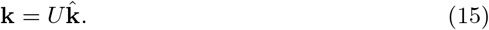

The convolution product on 𝒢 _L_ between **k** and **w** is the vector **k** * **w** such that

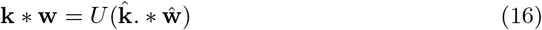

where ŵ is the graph Fourier transform of **w** defined as in (14). We choose *K*[**w**] as the convolution product of **w** with **k**:

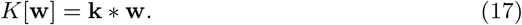

### 3.4 Model 4: spreading of *τ* via convolution on *𝒢* _nl_

In [33, 34], the authors introduce a graph convolution operator as a nonlocal model for the conversion from an healthy protein to a toxic one. Inspired by these works, we use their nonlocal operator to model the prion-like spreading of misfolded *τ* protein.

Let *M* ∈ ℝ^*N×N*^ be the matrix whose elements *m*_*i*,*j*_, *i, j* = 1, …, *N*, are the length of the shortest path from vertex *i* to vertex *j* in the graph 𝒢_NL_. If two vertices are not connected, *m*_*i*,*j*_ = 0. Moreover, let 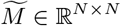 be the matrix with elements

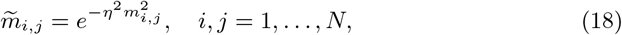

where *η* is a positive parameter. Finally, let *C* ∈ ℝ^*N×N*^ be the matrix defined as

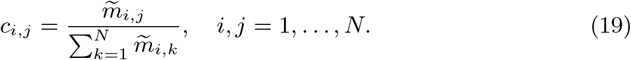

Then, the operator *K*[**w**] is chosen as

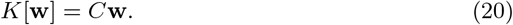

## 4 Methods

### 4.1 Data Availability

Data used in the preparation of this article were obtained from the ADNI Initiative (ADNI) database (https://adni.loni.usc.edu/). As described on the ADNI website, ADNI was launched in 2003 as a public-private partnership, led by Principal Investigator Michael W. Weiner, MD. The primary goal of ADNI has been to test whether serial MRI, positron emission tomography (PET), other biological markers, and clinical and neuropsychological assessment can be combined to measure the progression of mild cognitive impairment (MCI) and early Alzheimers disease. For up-to-date information, see www.adni-info.org.

### 4.2 Subjects

The study included a total of 261 participants: 238 Cognitive Normal (CN) subjects (mean age ±*st*.*dev*. = 72.7*y* ± 8.7*y*; *M* : *F* = 86 : 152), and 23 AD (mean age ±*st*.*dev*. = 73.1*y* ± 9.6*y*; *M* : *F* = 11 : 12) from the Alzheimers Disease Neuroimaging Initiative (ADNI). All data were downloaded from ADNI3. To quantitatively measure the concentration of *τ* in the brain regions, we considered subjects undergone a PET imaging with the radiotracer [^18^F]-AV1451 and a corresponding Accelerated sagittal T1-weighted anatomical image acquired using a 3D magnetization prepared rapid acquisition gradient echo (MPRAGE) sequence from Siemens 3T MRI scanners to minimize effects of inter-scanner variability. PET and MRI imaging was chosen to be acquired at no more than 3 months of temporal distance one from the other.

### 4.3 PET acquisition and processing

[^18^F]-AV1451 imaging acquisition and pre-processing in ADNI3 consists of the baseline [^18^F]-AV1451 PET scan (first scan acquired) for each subject, injection of 370 MBq (10 mCi) of tracer and 30 min dynamic scan consisting of six 5-min frames 75 min post-injection. Pre-processing consists of co-registration of separated frames to minimize patient motion effect; average of the 5 frames to obtain a single 30 min PET scan; re-orientering to a standardized space and normalization of intensity and finally the application of a filter specific for the type of scanner.

### 4.4 MRI structural imaging acquisition and processing

The MRI structural images chosen within 3 months to the date of [^18^F]-AV151 PET, were pre-processed, segmented and parcellated with FreeSurfer 6.0 (http://surfer.nmr.mgh.harvard.edu/), in order to subdivide the brain volumes into a set of 83 anatomical cortical and sub-cortical region of interest (ROIs), separated between left and right hemisphere (with the addition of the brain stem) and belonging to six main networks within the brain: the frontal, the parietal, the occipital, the temporal and the limbic lobes, plus the basal ganglia [13] [28]. The subdivision just described was chosen also based on the prior hypothesis that the limbic system is specially affected by the disease. A complete list of the regions, the abbreviation of their names and the network they belong to can be seen in the Appendix 7. Nodes of the brain graphs were put in correspondence with the 83 ROIs, considering the anatomical information of each vertex, to compare *τ* concentrations relative to the 83 ROIs in the PET images and the concentrations estimated by the mathematical models.

### 4.5 PET and MRI co-registration and quantification of *τ*

To calculate the differences in [^18^F]-AV1451 uptake between AD and CN we co-registered the PET images of each subject to the corresponding MRI structural image using PETSurfer (https://surfer.nmr.mgh.harvard.edu/fswiki/PetSurfer). Once the PET images are aligned in the same space of the structural image, since the ROIs parcellization is in the latter space, we can calculate the *τ* concentration in each of the 83 ROIs. To have an absolute quantification of each subject we normalized the concentration values to the cerebellum value, since it is considered a reference region for [^18^F]-AV1451 metabolism.

### 4.6 Statistical analysis

Firstly, the normality and homogeneity of variances of the distributions of *τ* concentrations for each group (AD and CN) were tested using the Shapiro-Wilks and Levenes tests. Then the distribution of [^18^F]-AV1451 PET uptake between AD and CN was calculated by a Mann-Whitney-Wilcoxon Test. The null hypothesis was rejected when p-values were below 0.05. In these cases, the post hoc test, Tukeys HSD test, was employed. Post hoc tests control the family-wise error rate by the Benjamini-Hochberg step-up procedure. The ROIs for which the null hypothesis was rejected were considered significantly different between AD and CN. Significant ROIs were ordered from the most significantly different between AD and CN to the less significant until the non-significant ROIs and the corresponding value of [^18^F]-AV1451 concentration was considered. In Figure 1 we depict the 83 cortical and subcortical nodes of the brain with colors corresponding to different *τ* concentration.

**Figure 1:**
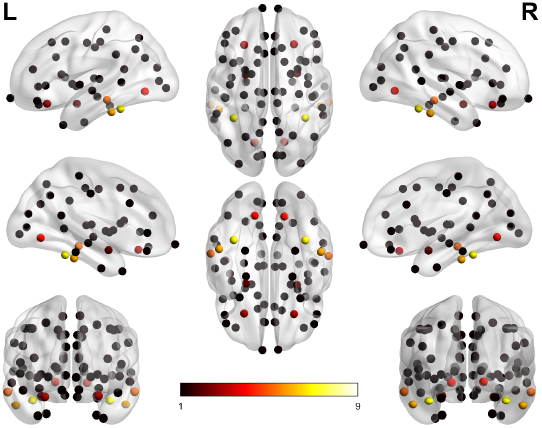
Nodes of the 83 cortical and subcortical ROIs of the brain. Color defines the nodes in which the concentration of *τ* is significantly greater in AD with respect to CN. The color-code defines the increasing order in *τ* concentration.

For the statistical analysis, we used Python libraries.

## 5 Results

This section presents the results of the numerical simulations of the four mathematical models corresponding to the different choices for the operator *K* governing *τ* spreading. All simulations were conducted using Matlab R2021a on an Intel Core i5 processor with 2.50GHZ and Windows operating system. The codes used for the current experiments can be made available upon request to the authors.

### 5.1 Experimental setting

The differential system (9) has been numerically solved by using the Matlab function ode45 implementing the Dormand-Prince 5(4) Runge-Kutta method with a variable time step for efficient computation [11]. Since misfolded *τ* protein has been shown to originate in the entorhinal region of the brain, the source term *s*_**w**_ (*t*) has been chosen non-null and equal to one only on those vertices belonging to the entorhinal region, i.e. for all *t >* 0 we have:

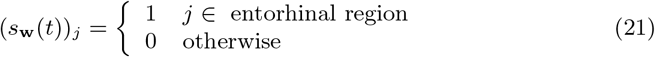

where (*s*_**w**_ (*t*))_*j*_ denotes the *j*th component of *s*_**w**_ (*t*). The seeding region for misfolded *τ* is shown in Figure 2.

**Figure 2:**
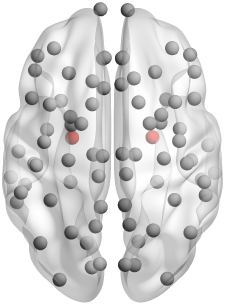
Nodes of the 83 ROIs corresponding to the cortical and subcortical regions of the brain. Color nodes correspond to the enthorinal regions.

The differential system has been integrated from the initial time *t*_0_ = 0 to the final time *t*_*f*_ = 150. The final time was chosen to correspond to the stabilisation of the solutions. In the numerical simulations of the four models, we have fixed the values of all parameters of system (9), except *γ*_3_, as reported in Table 1. These values have been chosen, on the basis of different tests, as the ones giving better results compared with clinical data.

**Table 1:** Fixed parameter values for system (9).

| $\gamma_1$ | $\gamma_2$ | $\alpha$ | $C_{\mathbf{u}_1}$ | $C_{\mathbf{w}}$ | $\sigma_1$ | $\sigma_2$ | $\sigma_3$ | $\sigma_4$ |
| --- | --- | --- | --- | --- | --- | --- | --- | --- |
| 0.001 | 0.001 | 0.12 | 0.1 | 0.5 | 0.1 | 0.1 | 0.1 | 0.11 |

Moreover we have fixed for the four models, with the same criterion as above, the parameters reported in Table 2. The procedure used for identifying, for each model, a good *γ*_3_ value will be described in the following subsection.

**Table 2:** Fixed parameter values for the four models.

| $R_p$ | $\delta$ | $\tilde{\delta}$ | $\eta$ |
| --- | --- | --- | --- |
| 25 | 30 | 15 | 5 |

## 5.2 Identification of the parameter *γ*_3_

The statistical analysis described in Section 4.6 shows that the distribution of ^18^F-AV1451 uptake is not normal for all ROIs. Subsequently, the non-parametric Mann-Whitney-Wilcoxon Test with a Bonferroni correction for multiple comparisons showed that 29 ROIs significantly differ between AD and CN. Table 3 shows the six ROIs which are mostly different (*p*-value¡ 10^*−*5^) and their respective avaraged *τ* concentrations. The most significant ROIs for the accumulation of *τ* in AD with respect to CN are, in decreasing order, the inferior temporal region, the fusiform region, the middle temporal region, the lateral orbitofrontal region, the amygdala and the lingual region. These six regions belong to three different networks (see Appendix 7): the inferior temporal and middle temporal regions belong to the temporal network, the fusiform and the lingual regions are part of the occipital network, and the lateral orbitofrontal region and the amygdala are included in the limbic network. Since the brain is organized in several functional discrete networks, as introduced in Section 4, it is convenient to refer to the networks to which these regions belong rather than to the single ROIs. For this reason, we average the values of *τ* protein in the significant regions belonging to the same network. Let 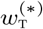 be the average of the *τ* values obtained from clinical data in the inferior temporal and middle temporal regions. Analogously, let 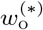 be the mean *τ* value in the fusiform and the lingual regions and let 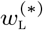 be the mean *τ* value in the later orbital and the amygdala regions.

**Table 3:**
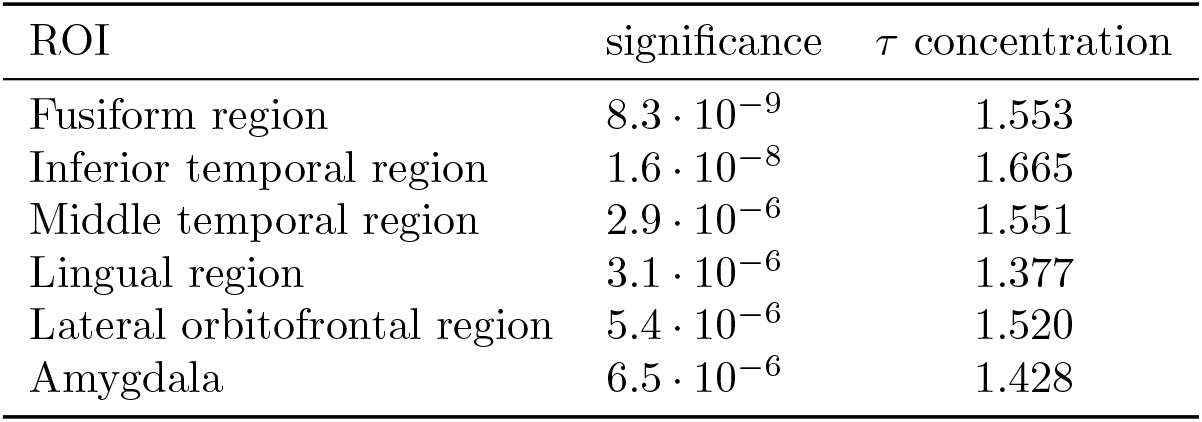
ROIs with a *τ* concentration significantly different between AD and CN and *τ* concentration in AD.

To evaluate the results obtained from four models, we also consider the value of *τ* protein in the sensorimotor network that is known not to be characterized by pathological accumulation of *τ* in AD. A good model for AD progression should, in fact, describe the evolution of *τ* both in the brain areas affected by the disease and in those that are not influenced. Therefore, let 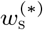 be the average *τ* value in the paracentral, postcentral, precentral and superior frontal regions forming the sensorimotor network. The *τ* mean values obtained from clinical data constitute our benchmark; they are ordered in decreasing order as follows

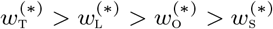

resulting in the string

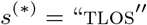

which therefore represents the decreasing order of *τ* values in clinical data. Brain regions with a higher concentration of misfolded *τ* are more deteriorated; therefore, *s*^(*\**)^ represents a *clinical deterioration pattern* of a brain with AD. We stress that this deterioration pattern considers only the brain regions that are significative for AD, according to our statistical analysis. Mathematical models should reproduce in the best possible way this deterioration pattern to be reliable tools for the analysis of AD progression. To evaluate the ability of the models to reproduce the deterioration pattern, we consider for each model *j, j* = 1, …, 4, the values of *τ* protein at the final time *t*_*f*_ . We average the values on the vertices belonging to the inferior temporal and middle temporal regions to get 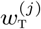, *j* = 1, …, 4. Proceeding similarly, we obtain 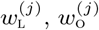 and 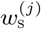 as mean *τ* values on the vertices belonging to the later orbital and the amygdala regions, the fusiform and the lingual regions and the paracentral, postcentral, precentral and superior frontal regions, respectively. For each model *j*, we define the string *s*^(*j*)^ formed using the letters T, L, O, S sorted in decreasing order of *τ* values 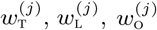 and 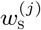 . We use the Hamming Distance (HD) between the strings *s*^(*\**)^ and *s*^(*j*)^, *j* = 1, …, 4 to measure the distance between the clinical and the models’ deterioration patterns. We recall that the Hamming distance between two strings is the number of positions at which the two strings differ [23]. The value of the parameter *γ*_3_ has been experimentally determined by trial and error to identify, for each model, the best outcome in terms of Hamming distance. Since, for each model, intervals of values exist for *γ*_3_ giving the same optimal Hamming distance value, we identify the best *γ*_3_ value in such intervals as follows. We define the vector

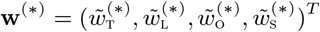

whose components, that are normalized *τ* values ranging from 0 to 1, are defined as follows:

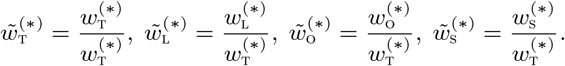

Then, we define the vectors

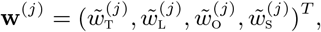

where

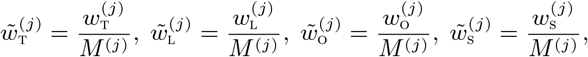

and 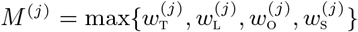. As a measure of the distance between **w**^(*\**)^ and **w**^(*j*)^ we can use the Root Mean Square Error (RMSE) and the Mean Absolute Error (MAE). We recall that, given a computed vector **w** ∈ ℝ^4^ of *τ* values, these error measures are respectively defined as

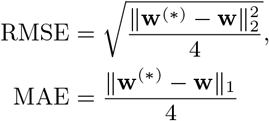

where ‖·‖_2_ and ‖·‖_2_ denote the *L*_2_ and *L*_1_ norms. Among all the *γ*_3_ values corresponding to the same optimal HD value, we heuristically determine the value which achieves the best RMSE reduction. This leads to four values for *γ*_3_, one for each model, shown in Table 4. Observe that the identified optimal *γ*_3_ value is the same for Model 1 and Model 2.

**Table 4:** Identified optimal values for *γ*_3_.

| Model 1 | Model 2 | Model 3 | Model 4 |
| --- | --- | --- | --- |
| 0.001 | 0.001 | 150 | 0.05 |

### 5.3 Numerical results

Table 5 shows the clinical and computed deterioration patterns; Tables 6 reports the corresponding HD, RMSE and MAE values.

**Table 5:** Clinical and models’ deterioration patterns.

| Clinical Data | Model 1 | Model 2 | Model 3 | Model 4 |
| --- | --- | --- | --- | --- |
| T | L | T | L | L |
| L | T | L | S | S |
| O | O | O | T | O |
| S | S | S | O | T |

**Table 6:** Numerical results for the four models.

|  | Model 1 | Model 2 | Model 3 | Model 4 |
| --- | --- | --- | --- | --- |
| HD | 2 | 0 | 4 | 3 |
| RMSE | 0.11574 | 0.11543 | 0.10931 | 0.10615 |
| MAE | 0.09256 | 0.09212 | 0.08911 | 0.08950 |

In Figure 3, we show the deterioration pattern in the significative brain regions from clinical data and from the four different models.

**Figure 3:**
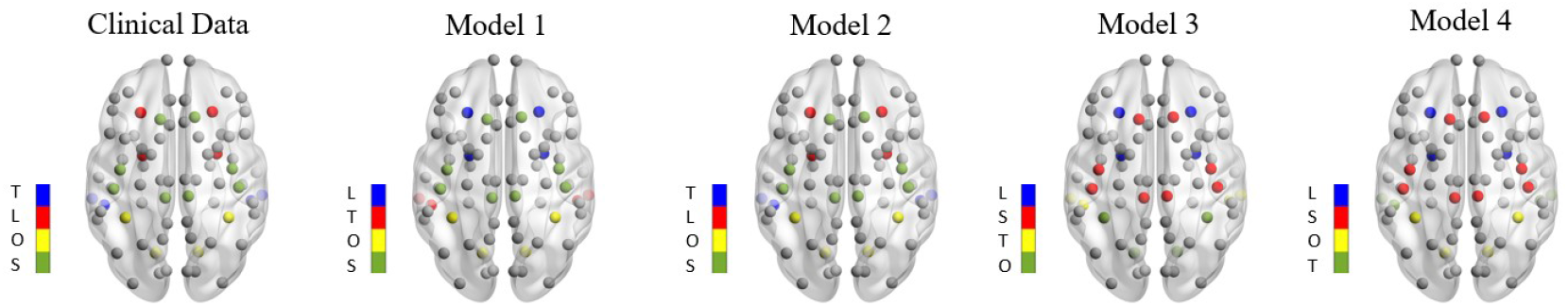
Deterioration pattern in the significative brain regions. From left to right: clinical data, Model 1, Model 2, Model 3 and Model 4. The non-significative regions are displayed in grey. In the colorbar, the colors correspond to decreasing *τ* values in the significative regions.

From the results shown above, it is clear that the only model capable of reproducing the clinical deterioration pattern is Model 2. A diffusion operator along the cumulative connectome seems to be the most appropriate operator for modelling the spreading of *τ* protein in a brain with AD. Model 1 uses a diffusion operator too, but comparing the deterioration patterns of Model 1 and Model 2, it is evident that the cumulative connectome, encoding information on brain regions connected along long distances, is a key tool. Even if Model 1 does not exactly reproduce the clinical deterioration pattern, it is able to distinguish between deteriorated and non-deteriorated regions, since the letter S is in the last position in the string *w*^(1)^, as it should be. We observe from Table 6 that models 3 and 4 give the best results in terms of RMSE and MAE reduction. However, they do not reproduce the deterioration pattern of AD and do not distinguish between deteriorated and non-deteriorated regions (the letter S is in the second position both in *w*^(3)^ and *w*^(4)^). For this reason, they do not seem to be good models for the spreading of misfolded *τ* in presence of AD.

## 6 Discussion

In this study we had two main objectives. We were interested in: 1) comparing different modellistic possibilities concerning the spreading of *τ* protein in a brain with AD, when also *Aβ* protein is present and the synergy between the two is considered; 2) producing clinical data which can be compared with the outputs of the models in order to verify their reliability. We believe that the lack of comparison between the results of a theoretical model and clinical data could be considered a deficiency that should be rectified in order to support the usefulness of mathematical models in AD [30]. To this end, we evolved the two proteins on appropriate networks, created from medical data of human connectomes. The need for different connectomes was dictated by both the physiology of the brain and the biological characteristics of the proteins themselves. We therefore considered for the evolution of *Aβ*, which travels only on short distances, an intrinsic proximity connectome, and a standard diffusion Laplacian on it. With regard to *τ* spreading, which is supposed to possibly travel with a prion-like mechanism and over long distances, we considered four distinct mathematical models on as many networks: a diffusion via Laplacian on a structural connectome and on a cumulative connectome, a spreading via convolution on two further different connectomes. We emphasize that the so-called intrinsic proximity connectome and the cumulative connectome have been introduced in this work for the first time. We identified from public multimodal data, with a careful statistical analysis, six regions relevant when it comes to AD and we evaluated *τ* concentrations in them, obtaining a degradation pattern that was crucial for us to verify the goodness of the models.

By comparison of the results of the simulations with clinical data, we saw that only the model using the cumulative connectome was able to reproduce correctly the clinical degradation pattern, that also includes a network that is not damaged by the disease. We hypothesize that this result is due to the fact that the cumulative connectome encodes information on brain regions connected along long distances. On the basis of these findings we feel we can say that the models are not all equivalent, and that the comparison with clinical data is a diriment element to be able to assess their reliability. Of course, the results obtained here relate to the specific case of AD, and it is plausible that for other diseases the same models give different performances. As a next step we are going to apply the same paradigm of analysis, i.e. close comparison with appropriate clinical data, to the models enriched by taking into account atrophy, to be more realistic and see if something relevant happens. We believe that a highly interdisciplinary study, such as the one reported in the paper, could be a very promising direction in which cutting-edge biomedical mathematical research could be heading.

## Author Contributions

All authors have contributed equally to this work. All authors have read and agreed to the published version of the manuscript.

## Funding

This research was supported by the AlmaIdea 2022 programme of the Alma Mater Studiorum - Universit di Bologna, project n. CUPJ45F21002000001.

## Institutional Review Board Statement

As per ADNI protocols, all procedures performed in studies involving human participants were in accordance with the ethical standards of the institutional and/or national research committee and with the 1964 Helsinki declaration and its later amendments or comparable ethical standards. More details can be found at adni.loni.usc.edu. (This article does not contain any studies with human participants performed by any of the authors)

## Informed Consent Statement

Authors received the consent of publication from ADNI.

## Data Availability Statement

Publicly available datasets were analyzed in this study. This data can be found at these urls: http://adni.loni.usc.edu and https://braingraph.org see [42, 43].

All data produced by the authors are available upon request from the authors.

## Acknowledgments

Data collection and sharing for this project was funded by the Alzheimer’s Disease Neuroimaging Initiative (ADNI) (National Institutes of Health Grant U01 AG024904) and DOD ADNI (Department of Defense award number W81XWH-12-2-0012). ADNI is fundedby the National Institute on Aging, the National Institute of Biomedical Imaging and Bioengineering, and through generous contributions from the following: AbbVie, Alzheimers Association; Alzheimers Drug Discovery Foundation; Araclon Biotech; BioClinica, Inc.; Biogen; Bristol-Myers Squibb Company; CereSpir, Inc.; Cogstate; Eisai Inc.; Elan Pharmaceuticals, Inc.; Eli Lilly and Company; EuroImmun; F. Hoffmann-La Roche Ltd and its affiliated company Genentech, Inc.; Fujirebio; GE Healthcare; IXICO Ltd.; Janssen Alzheimer Immunotherapy Research & Development, LLC.; Johnson & Johnson Pharmaceutical Research & Development LLC.; Lumosity; Lundbeck; Merck & Co., Inc.; Meso Scale Diagnostics, LLC.; NeuroRx Research; Neurotrack Technologies; Novartis Pharmaceuticals Corporation; Pfizer Inc.; Piramal Imaging; Servier; Takeda Pharmaceutical Company; and Transition Therapeutics. The Canadian Institutes of Health Research is providing funds to support ADNI clinical sites in Canada. Private sector contributions are facilitated by the Foundation for the National Institutes of Health (www.fnih.org). The grantee organization is the Northern California Institute for Research and Education, and the study is coordinated by the Alzheimers Therapeutic Research Institute at the University of Southern California. ADNI data are disseminated by the Laboratory for Neuro Imaging at the University of Southern California.

## Conflicts of Interest

The authors declare no conflicts of interest.

## 7 Appendix

**Table 7:** List of anatomical cortical and sub-cortical ROIs identified with FreeSurfer, and the six networks: frontal, occipital, temporal, limbic, sensorimotor network and basal ganglia.

| Region of interest | Network |
| --- | --- |
| Nucleus accumbens | Limbic |
| Amygdala | Limbic |
| Bankssts | Temporal |
| Brain Stem | Limbic |
| Caudal anterior cingulate gyrus | Limbic |
| Caudal middle frontal gyrus | Frontal |
| Caudate | Basal ganglia |
| Cuneus | Occipital |
| Entorhinal gyrus | Limbic |
| Frontal pole | Frontal |
| Fusiform gyrus | Occipital |
| Hippocampus | Limbic |
| Inferior parietal gyrus | Parietal |
| Inferior temporal gyrus | Temporal |
| Insula | Limbic |
| Isthmus cingulate gyrus | Limbic |
| Lateral occipital gyrus | Occipital |
| Lateral orbito frontal gyrus | Limbic |
| Lingual gyrus | Occipital |
| Medial orbito frontal gyrus | Limbic |
| Middle temporal gyrus | Temporal |
| Pallidum | Basal ganglia |
| Paracentral gyrus | Sensorimotor |
| Parahippocampal gyrus | Limbic |
| Pars opercularis | Frontal |
| Pars orbitalis | Frontal |
| Pars triangularis | Frontal |
| Peri calcarine gyrus | Occipital |
| Postcentral gyrus | Sensorimotor |
| Posterior cingulate gyrus | Limbic |
| Precentral gyrus | Sensorimotor |
| Precuneus | Parietal |
| Putamen | Basal ganglia |
| Rostral anterior cingulate gyrus | Limbic |
| Rostral middle frontal gyrus | Frontal |
| Superior frontal gyrus | Sensorimotor |
| Superior parietal gyrus | Parietal |
| Superior temporal gyrus | Temporal |
| Supramarginal gyrus | Parietal |
| Temporal pole | Temporal |
| Transverse temporal gyrus | Temporal |

## Notes

### Competing Interest Statement

The authors have declared no competing interest.

## References

[1] Zeshan Ahmed, Jane Cooper, Tracey K Murray, Katya Garn, Emily McNaughton, Hannah Clarke, Samira Parhizkar, Mark A Ward, Annalisa Cavallini, Samuel Jackson, et al. A novel in vivo model of tau propagation with rapid and progressive neurofibrillary tangle pathology: the pattern of spread is determined by connectivity, not proximity. Acta neuropathologica, 127:667–683, 2014.

[2] Danielle S Bassett and Edward T Bullmore. Human brain networks in health and disease. Current opinion in neurology, 22(4):340–347, 2009.

[3] Rachel E Bennett, Sarah L DeVos, Simon Dujardin, Bianca Corjuc, Rucha Gor, Jose Gonzalez, Allyson D Roe, Matthew P Frosch, Rose Pitstick, George A Carlson, et al. Enhanced tau aggregation in the presence of amyloid β. The American journal of pathology, 187(7):1601–1612, 2017.

[4] Michiel Bertsch, Bruno Franchi, Ashish Raj, and Maria Carla Tesi. Macroscopic modelling of alzheimers disease: difficulties and challenges. Brain Multiphysics, 2:100040, 2021.

[5] Michiel Bertsch, Bruno Franchi, Maria Carla Tesi, and Veronica Tora. The role of a β and tau proteins in alzheimers disease: a mathematical model on graphs. Journal of Mathematical Biology, 87(3):49, 2023.

[6] George S Bloom. Amyloid-β and tau: the trigger and bullet in alzheimer disease pathogenesis. JAMA neurology, 71(4):505–508, 2014.

[7] Heiko Braak and Eva Braak. Neuropathological stageing of alzheimer-related changes. Acta neuropathologica, 82(4):239–259, 1991.

[8] Heiko Braak and Kelly Del Tredici. Alzheimers pathogenesis: is there neuron-to-neuron propagation? Acta neuropathologica, 121:589–595, 2011.

[9] Marc Aurel Busche and Bradley T Hyman. Synergy between amyloid-β and tau in alzheimers disease. Nature neuroscience, 23(10):1183–1193, 2020.

[10] Marco Catani and Marsel Mesulam. What is a disconnection syndrome? 2008.

[11] John R Dormand and Peter J Prince. A family of embedded runge-kutta formulae. Journal of computational and applied mathematics, 6(1):19–26, 1980.

[12] Simon Dujardin and Bradley T Hyman. Tau prion-like propagation: state of the art and current challenges. Tau Biology, pages 305–325, 2020.

[13] Stefania Evangelisti, Claudia Testa, Lorenzo Ferri, Laura Ludovica Gramegna, David Neil Manners, Giovanni Rizzo, Daniel Remondini, Gastone Castellani, Ilaria Naldi, Francesca Bisulli, et al. Brain functional connectivity in sleep-related hypermotor epilepsy. NeuroImage: Clinical, 17:873–881, 2018.

[14] Sveva Fornari, Amelie Schäfer, Mathias Jucker, Alain Goriely, and Ellen Kuhl. Prion-like spreading of alzheimers disease within the brains connectome. Journal of the Royal Society Interface, 16(159):20190356, 2019.

[15] Sveva Fornari, Amelie Schäfer, Ellen Kuhl, and Alain Goriely. Spatially-extended nucleation-aggregation-fragmentation models for the dynamics of prion-like neurodegenerative protein-spreading in the brain and its connectome. Journal of theoretical biology, 486:110102, 2020.

[16] Maria Laura Giuffrida, Filippo Caraci, Bruno Pignataro, Sebastiano Cataldo, Paolo De Bona, Valeria Bruno, Gemma Molinaro, Giuseppe Pappalardo, Angela Messina, Angelo Palmigiano, et al. β-amyloid monomers are neuroprotective. Journal of Neuroscience, 29(34):10582–10587, 2009.

[17] Michel Goedert and Maria Grazia Spillantini. Propagation of tau aggregates. Molecular brain, 10:1–9, 2017.

[18] Alain Goriely, Ellen Kuhl, and Christian Bick. Neuronal oscillations on evolving networks: dynamics, damage, degradation, decline, dementia, and death. Physical review letters, 125(12):128102, 2020.

[19] Alexander Grigoryan. Introduction to analysis on graphs, volume 71. American Mathematical Soc., 2018.

[20] Michel J Grothe, Henryk Barthel, Jorge Sepulcre, Martin Dyrba, Osama Sabri, Stefan J Teipel, Alzheimer’s Disease Neuroimaging Initiative, and Alzheimer’s Disease Neuroimaging Initiative. In vivo staging of regional amyloid deposition. Neurology, 89(20):2031–2038, 2017.

[21] I Grundke-Iqbal, K Iqbal, YC Tung, M Quinlan, HM Wisniewski, and LI Binder. Abnormal phosphorylation of the microtubule-associated protein?(tau) in alzheimer cytoskeletal pathology. Alzheimer Disease & Associated Disorders, 1(3):202, 1987.

[22] Christian Haass and Dennis J Selkoe. Soluble protein oligomers in neurodegeneration: lessons from the alzheimer’s amyloid β-peptide. Nature reviews Molecular cell biology, 8(2):101–112, 2007.

[23] Kermit Hutcheson. A test for comparing diversities based on shannon formula. Journal of theoretical Biology, 29:151–154, 1970.

[24] Lars M Ittner and Jürgen Götz. Amyloid-β and taua toxic pas de deux in alzheimer’s disease. Nature Reviews Neuroscience, 12(2):67–72, 2011.

[25] JunHyun Kim, Minhong Jeong, Wesley R Stiles, and Hak Soo Choi. Neuroimaging modalities in alzheimers disease: diagnosis and clinical features. International journal of molecular sciences, 23(11):6079, 2022.

[26] Jennifer A McNab, Brian L Edlow, Thomas Witzel, Susie Y Huang, Himanshu Bhat, Keith Heberlein, Thorsten Feiweier, Kecheng Liu, Boris Keil, Julien Cohen-Adad, et al. The human connectome project and beyond: initial applications of 300 mt/m gradients. Neuroimage, 80:234–245, 2013.

[27] M-Marchsel Mesulam. A cortical network for directed attention and unilateral neglect. Annals of Neurology: Official Journal of the American Neurological Association and the Child Neurology Society, 10(4):309–325, 1981.

[28] M-Marsel Mesulam. From sensation to cognition. Brain: a journal of neurology, 121(6):1013–1052, 1998.

[29] Melanie Meyer-Luehmann, Tara L Spires-Jones, Claudia Prada, Monica Garcia-Alloza, Alix De Calignon, Anete Rozkalne, Jessica Koenigsknecht-Talboo, David M Holtzman, Brian J Bacskai, and Bradley T Hyman. Rapid appearance and local toxicity of amyloid-β plaques in a mouse model of alzheimers disease. Nature, 451(7179):720–724, 2008.

[30] Seyedadel Moravveji, Nicolas Doyon, Javad Mashreghi, and Simon Duchesne. A scoping review of mathematical models covering alzheimer’s disease progression. Frontiers in Neuroinformatics, 18:1281656, 2024.

[31] Kenjiro Ono, Margaret M Condron, and David B Teplow. Structure–neurotoxicity relationships of amyloid β-protein oligomers. Proceedings of the National Academy of Sciences, 106(35):14745–14750, 2009.

[32] Rik Ossenkoppele, Gil D Rabinovici, Ruben Smith, Hanna Cho, Michael Schöll, Olof Strandberg, Sebastian Palmqvist, Niklas Mattsson, Shorena Janelidze, Alexander Santillo, et al. Discriminative accuracy of [18f] flortaucipir positron emission tomography for alzheimer disease vs other neurodegenerative disorders. Jama, 320(11):1151–1162, 2018.

[33] S Pal and R Melnik. Nonlocal multiscale interactions in brain neurodegenerative protein dynamics and coupled proteopathic processes. In Proceedings of the 9th Edition of the International Conference on Computational Methods for Coupled Problems in Science and Engineering (Coupled Problems 2021), Online Event, pages 14–16, 2021.

[34] Swadesh Pal and Roderick Melnik. Nonlocal models in the analysis of brain neurodegenerative protein dynamics with application to alzheimers disease. Scientific Reports, 12(1):7328, 2022.

[35] Ashish Raj. Graph models of pathology spread in alzheimer’s disease: an alternative to conventional graph theoretic analysis. Brain connectivity, 11(10):799–814, 2021.

[36] Ashish Raj, Amy Kuceyeski, and Michael Weiner. A network diffusion model of disease progression in dementia. Neuron, 73(6):1204–1215, 2012.

[37] Ashish Raj, Veronica Tora, Xiao Gao, Hanna Cho, Jae Yong Choi, Young Hoon Ryu, Chul Hyoung Lyoo, and Bruno Franchi. Combined model of aggregation and network diffusion recapitulates alzheimer’s regional tau-positron emission tomography. Brain connectivity, 11(8):624–638, 2021.

[38] Roberta Ricciarelli and Ernesto Fedele. The amyloid cascade hypothesis in alzheimer’s disease: it’s time to change our mind. Current neuropharmacology, 15(6):926–935, 2017.

[39] David I Shuman, Sunil K Narang, Pascal Frossard, Antonio Ortega, and Pierre Vandergheynst. The emerging field of signal processing on graphs: Extending high-dimensional data analysis to networks and other irregular domains. IEEE signal processing magazine, 30(3):83–98, 2013.

[40] Scott A Small and Karen Duff. Linking aβ and tau in late-onset alzheimer’s disease: a dual pathway hypothesis. Neuron, 60(4):534–542, 2008.

[41] Olaf Sporns and Jonathan D Zwi. The small world of the cerebral cortex. Neuroinformatics, 2:145–162, 2004.

[42] Balazs Szalkai, Csaba Kerepesi, Balint Varga, and Vince Grolmusz. Parameterizable consensus connectomes from the human connectome project: the budapest reference connectome server v3. 0. Cognitive neurodynamics, 11:113–116, 2017.

[43] Balazs Szalkai, Csaba Kerepesi, Balint Varga, and Vince Grolmusz. Highresolution directed human connectomes and the consensus connectome dynamics. PLoS One, 14(4):e0215473, 2019.

[44] OG Tatarnikova, MA Orlov, and NV Bobkova. Beta-amyloid and tau-protein: Structure, interaction, and prion-like properties. Biochemistry (Moscow), 80:1800–1819, 2015.

[45] Travis B Thompson, Pavanjit Chaggar, Ellen Kuhl, Alain Goriely, and Alzheimers Disease Neuroimaging Initiative. Protein-protein interactions in neurodegenerative diseases: A conspiracy theory. PLoS computational biology, 16(10):e1008267, 2020.

[46] Justin Torok, Chaitali Anand, Parul Verma, and Ashish Raj. Connectome-based biophysics models of alzheimers disease diagnosis and prognosis. Translational Research, 254:13–23, 2023.

